# Parental legacy, demography, and introgression influenced the evolution of the two subgenomes of the tetraploid *Capsella bursa-pastoris* (Brassicaceae)

**DOI:** 10.1101/234096

**Authors:** Dmytro Kryvokhyzha, Adriana Salcedo, Mimmi C. Eriksson, Tianlin Duan, Nilesh Tawari, Jun Chen, Maria Guerrina, Julia M. Kreiner, Tyler V. Kent, Ulf Lagercrantz, John R. Stinchcombe, Sylvain Glémin, Stephen I. Wright, Martin Lascoux

## Abstract

Allopolyploidy is generally perceived as a major source of evolutionary novelties and as an instantaneous way to create isolation barriers. However, we do not have a clear understanding of how two subgenomes evolve and interact once they have fused in an allopolyploid species and how isolated they are from their relatives. Here, we address these questions by analyzing genomic and transcriptomic data of allotetraploid *Capsella bursa-pastoris* in three differentiated populations, Asia, Europe and the Middle East. We phased the two subgenomes, one descended from the outcrossing and highly diverse *Capsella grandiflora* (Cg) and the other one from the selfing and genetically depauperate *Capsella orientalis* (Co). For each subgenome, we assessed its relationship with the diploid relatives, temporal change of effective population size *N_e_*, signatures of positive and negative selection, and gene expression patterns. Introgression between *C. bursa-pastoris* and its diploid relatives was widespread and the two subgenomes were impacted differentially depending on geographic region. In all three regions, Ne of the two subgenomes decreased gradually and the Co subgenome accumulated more deleterious changes than Cg. Selective sweeps were more common on the Cg subgenome in Europe and the Middle East, and on the Co subgenome in Asia. In contrast, differences in expression were limited with the Cg subgenome slightly more expressed than Co in Europe and the Middle-East. In summary, after more than 100,000 generations of co-existence, the two subgenomes of *C. bursa-pastoris* still retained a strong signature of parental legacy and were differentially affected by introgression and selection.

## Introduction

Allopolyploidy, the origin of polyploids from two different ancestral lineages, poses serious evolutionary challenges since the presence of two divergent sub-genomes may lead to perturbation of meiosis, conflicts in gene expression regulation, protein-protein interactions and/or transposable element suppression (1, 2). Whole genome duplication also masks new recessive mutations thereby decreasing selection efficacy (3, 4). This relaxation of selection, together with the strong speciation bottleneck and shift to self-fertilization that often accompany polyploidy (5), ultimately increases the frequency of deleterious mutations retained in the genome (6, 7). All of these consequences of allopolyploidy can have a negative impact on fitness and over evolutionary time may contribute to the patterns of duplicate gene loss, a process referred to as diploidization (4, 8, 9). Yet, allopolyploid lineages often not only establish and persist but may even thrive and become more successful than their diploid progenitors and competitors, with larger ranges and higher competitive ability (10–19). The success of allopolyploids is usually explained by their greater evolutionary potential. Having inherited two genomes that evolved separately, and sometimes under drastically different conditions, allopolyploids should have an increased genetic toolbox, assuming that the two genomes do not experience severe conflicts. This greater evolutionary potential of allopolyploids can be further enhanced by genomic rearrangements, alteration of gene expression and epigenetic changes (3, 4, 20–26).

All of these specific features come into play during the demographic history of allopolyploids. Demographic processes occurring when a species extends its range, such as successive bottlenecks or periods of rapid population growth in the absence of competition, are expected to have a profound impact on evolutionary processes, especially in populations at the front of the expansion range. Species that went through repeated bottlenecks during their range expansion are expected to have reduced genetic variation and higher genetic load than more ancient central populations (27, 28). Similarly, range expansions can also lead to contact and gene flow with introgression from related species. Such gene flow can in turn shift the evolutionary path of the focal species. Finally, range expansion will expose newly formed allopoly-ploid populations to divergent selective pressures, providing the possibility of differentially exploiting duplicated genes, and creating asymmetrical patterns of adaptive evolution in different parts of the range.

In this paper, we aim to characterize the evolution of the genome of a recent allopolyploid species during its range expansion. In particular, we explore whether the two subgenomes have similar or different evolutionary trajectories in term of hybridization, selection and gene expression. The widespread allopolyploid *C. bursa-pastoris* is a promising system for studying the evolution of polyploidy, with available information on its two progenitor diploid species and their current distribution. *C. bursa-pastoris*, a selfing species, originated from the hybridization of the *Capsella orientalis* and *Capsella grandiflora / rubella* lineages some 100-300 kya (9). *C. orientalis* is a genetically depauperate selfer occurring across the steppes of Central Asia and Eastern Europe. In contrast, *C. grandiflora* is an extremely genetically diverse obligate outcrosser which is primarily confined to a tiny distribution range in the mountains of Northern Greece and Albania. The fourth relative, *C. rubella*, a selfer recently derived from *C. grandiflora*, occurs around the Mediterranean Sea (Fig. 1A). There is evidence for unidirectional gene flow from *C. rubella* to *C. bursa-pastoris* (29). Among all *Capsella* species, only *C. bursa-pastoris* has a worldwide distribution (30), some of which might be due to extremely recent colonization and associated with human population movements (31). A recent study reveals that in Eurasia, *C. bursa-pastoris* is divided into three genetic clusters - Middle East, Europe and Asia - with low gene flow among them and strong differentiation both at the nucleotide and gene expression levels (31, 32). Reconstruction of the colonization history using unphased genomic data suggested that *C. bursa-pastoris* spread from the Middle East towards Europe and then into Eastern Asia. This colonization history resulted in a typical reduction of nucleotide diversity with the lowest diversity being in the most distant Asian population (31).

**Fig. 1.**
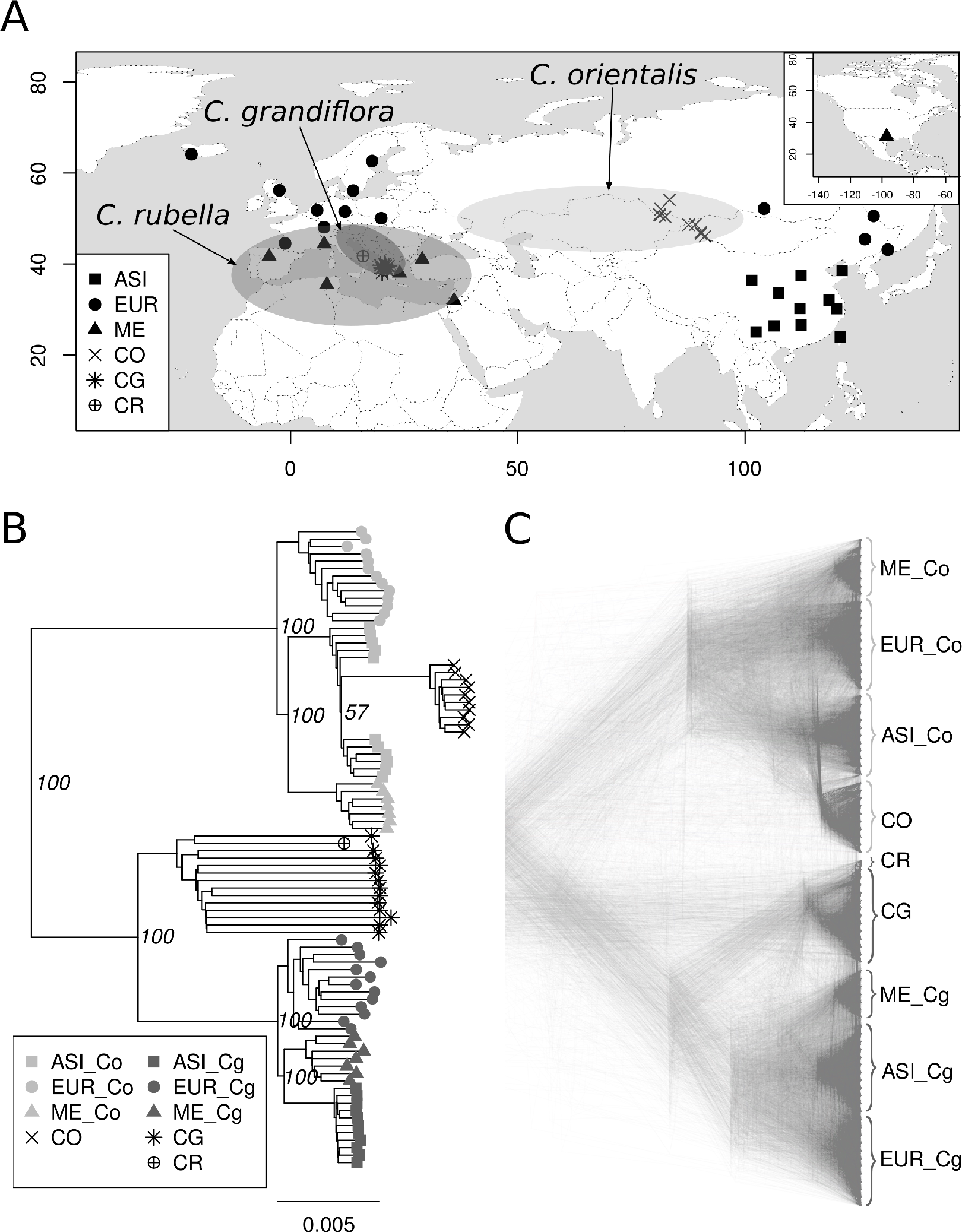
Distribution ranges, sampling locations and phylogenetic relationships of *Capsella* species used in this study. **A.** Approximative distribution ranges of *C. orientalis, C. grandiflora*, and *C. rubella* and sampling locations of *C. bursa-pastoris. C. bursa-pastoris* has a worldwide distribution, so its distribution range is not specifically depicted. ASI, EUR ME, CO, CG, CR indicate Asian, European and Middle Eastern populations of *C. bursa-pastoris, C. orientalis*, *C. grandiflora*, and *C. rubella*, respectively. The map is modified from Hurka et al. (30). **B.** Whole genome NJ tree showing the absolute divergence between different populations of *C. bursa-pastoris* at the level of subgenomes. The Co and Cg subgenomes are marked with corresponding names. The bootstrap support based on 100 replicates is shown only for the major clades. The root *N. paniculata* is not shown. **C.** Density tree visualizing of 1002 NJ trees reconstructed with 100 Kb sliding windows.

How the two distinct non-recombining subgenomes of *C. bursa-pastoris* contributed to its rapid population expansion and how they were in return affected by it, remains unclear. Previous studies either ignored the population history of *C. bursa-pastoris* or failed to consider the two subgenomes separately. In a recent study that does not consider the population demographic history within *C. bursa-pastoris*, Douglas et al.(9) concluded that there is no strong sign of diploidization in *C. bursa-pastoris* and most of its variation is the result of the legacy from the parental lineages with some relaxation of purifying selection caused by both the transition to self-fertilization and the greater masking of deleterious mutations. Kryvokhyzha et al. (32) considered population history but did not separate the two subgenomes, and showed that variation in gene expression among Asian, European and Middle Eastern accessions strongly reflects the population history with most of the differences among populations explained by genetic drift. We extend these previous studies by analyzing the genome-wide expression and polymorphism patterns of the two subgenomes of *C. bursa-pastoris* in 31 accessions sampled across its natural range in Eurasia. We demonstrate that the two subgenomes follow distinct evolutionary trajectories in different populations and that these trajectories are influenced by both range expansion and introgression from relatives. Our study illustrates the need to account for demographic and ecological differences among populations when studying the evolution of subgenomes of allopolyploid species.

## Results

**Phasing subgenomes.** The disomic inheritance of *C. bursa-pastoris* allowed us to successfully phase most of the heterozygous sites in the 31 samples analyzed in this study (Fig. 1A, Table S1). Out of 7.1 million high confidence SNPs, our phasing procedure produced an alignment of 5.4 million phased polymorphic sites across the 31 accessions of *C. bursa-pastoris*. Scaling these phased SNPs to the whole genome resulted in the alignment of 80.6 Mb that had the same level of heterozygosity as the unphased data. The alignment of these whole genome sequences of *C. bursapastoris* with 13 sequences of *C. grandiflora*, 10 sequences of *C. orientalis*, one sequence of *C. rubella* (the reference), and one sequence of *N. paniculata* used here as an outgroup, yielded 13 million polymorphic sites that we used in all analyses. The information for each accession is provided in Table S1.

To assess the quality of the phasing results, we constructed a phylogeny from the phased data. The separation of the two subgenomes was strongly supported in the reconstructed whole genome tree (Fig. 1B). The tree consisted of two highly supported (100% bootstrap) major clades grouping *C. grandiflora* and the *C. grandiflora / rubella* lineage descended subgenome of *C. bursa-pastoris* (hereafter the Cg subgenome), on the one hand, and *C. orientalis* and the *C. orientalis* lineage descended subgenome of *C. bursapastoris* (hereafter the Co subgenome), on the other hand. We also analyzed phylogenetic signals at a finer genomic scale using a sliding window approach with 100-kb window size (Fig. 1C). Exclusive monophyly of *C. orientalis* with the Co subgenome, and *C. grandiflora* and *C. rubella* with Cg subgenome was detected in 95% and 83% of trees, respectively (Fig. S1).

**Polymorphism and population structure of the two subgenomes.** For both subgenomes the three *C. bursapastoris* populations, Asia (ASI), Europe (EUR) and Middle East (ME), constituted well-defined phylogenetic clusters (Fig. 1B,C). However, the relationships of each subgenome with its parental species differed. The Cg subgenome formed a monophyletic clade with *C. grandiflora* at its base. In contrast, the Co subgenome was paraphyletic with *C. orientalis* that clustered within the ASI group instead of being outside of all *C. bursa-pastoris* Co subgenomes. This clustering was unexpected and suggested potential gene flow between the ASI group and *C. orientalis* or multiple origins of the Co subgenome. Nucleotide diversity was higher on the Cg subgenome than on the Co subgenome for both EUR and ME (Fig. 2, Table S2), though the difference was significant only for EUR (p-values: 0.005 and 0.154 for EUR and ME, respectively). The opposite pattern was observed for ASI (Fig. 2): there the nucleotide diversity in the Co subgenome was significantly higher than in the Cg subgenome (p-value < 0.0001). Interestingly, the diversity of the Co subgenome in all populations was significantly higher than the diversity of its parental species, *C. orientalis* (p-value < 0.0001).

**Fig. 2.**
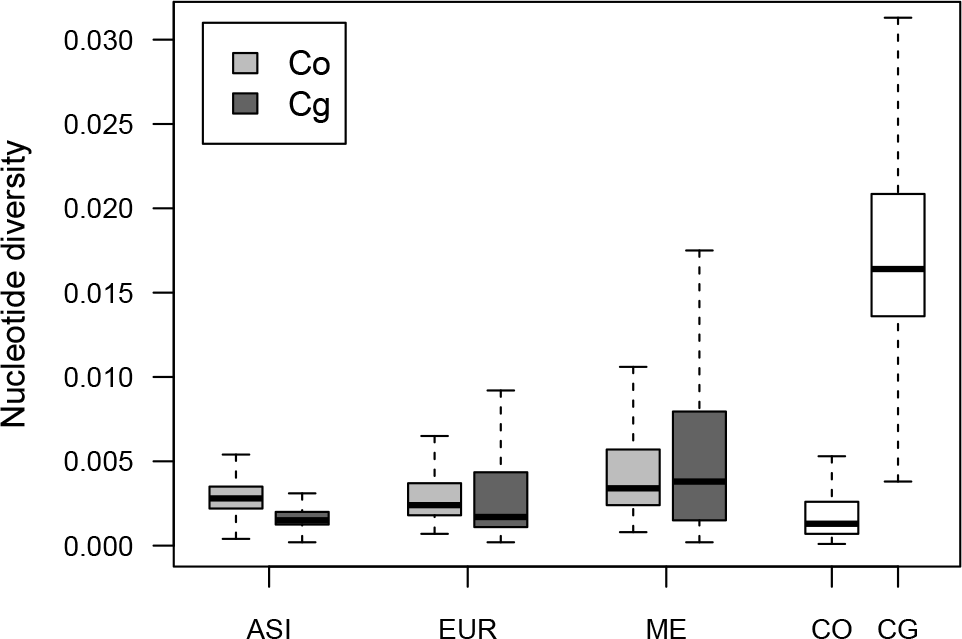
Variation in nucleotide diversity (*π*) between populations of *C. bursa-pastoris* and parental species. This boxplot shows *π* estimated along the genome using 100 Kb sliding windows. Co and Cg indicate *C, orientalis* and *C, grandiflora /rubella* descendant subgenomes, respectively. ASI, EUR ME, CO and CG correspond to Asian, European and Middle Eastern populations of *C, bursa-pastoris, C, orientalis*, and *C, grandiflora*, respectively.

**Temporal change in effective population size.** To reconstruct the changes in effective population size (*N_e_*) over time in the three *C. bursa-pastoris* populations and the two ancestral species, we used a pairwise sequentially Markovian coalescent model (PSMC). First, we reconstructed the demographic histories of *C. orientalis* and *C. grandiflora* (Fig. 3). In *C. grandiflora*, *N_e_*, was mostly constant with some slight decrease in the recent past, but the *N_e_* of *C. orientalis* decreased continuously. In *C. bursa-pastoris*, despite a simultaneous rapid range expansion, *N_e_* of EUR and ME populations also gradually decreased starting from around 100-200 kya. The ASI population showed a similar pattern but with population size recovery in the range 5-10 kya and a subsequent decrease to the same *N_e_* as in EUR and ME. The *N_e_* patterns of the two subgenomes were similar within each population. Overall, the *N_e_* history of *C. bursa-pastoris* was most similar to that of its selfing ancestor, *C. orientalis*. We also verified these PSMC results with SMC++, which can consider more than two haploid genomes and incorporates linkage disequilibrium (LD) in coalescent hidden Markov models (33). The general trend was globally the same but the recent decline of *C. orientalis* was sharper and fluctuations in *N_e_*, were more pronounced (Fig. S2). In summary, the overall pattern of *N_e_* change over time was mostly the same between the two subgenomes and between the three populations of *C. bursa-pastoris* and it was largely similar to the pattern observed for the diploid selfer *C. orientalis*.

**Fig. 3.**
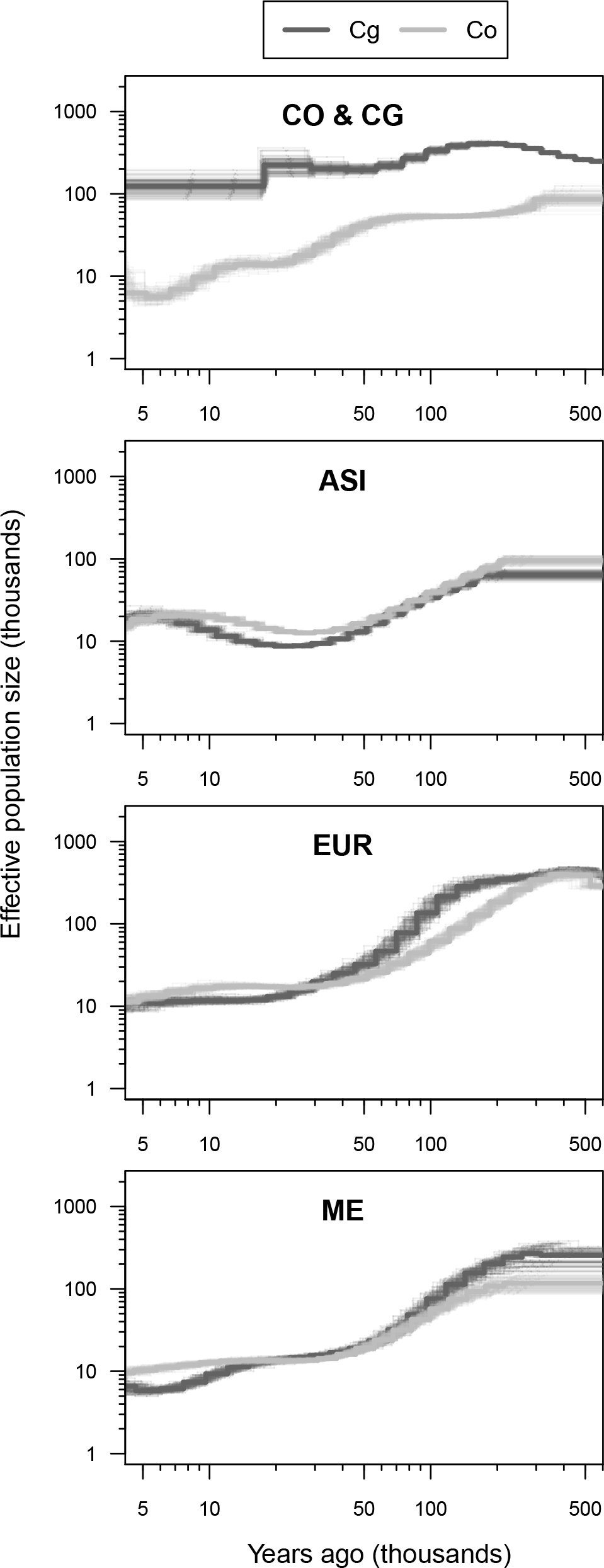
Population size histories of *C. bursa-pastoris* and its parental species. Effective population sizes were inferred with PSMC using whole-genome sequences from a pair of haplotypes per population (thick lines) and 100 bootstrap replicates (thin lines). The estimates for different pairs were similar and shown in the Supp. (Fig. S12). Co and Cg specify subgenomes of *C, bursa-pastoris* and corresponding parental species in the CO & CG plot. ASI, EUR, ME, CO & CG indicate Asian, European and Middle Eastern populations of *C, bursa-pastoris*, and *C, orientalis* and *C, grandiflora*, respectively. The axis are in log scale and the most recent times where PSMC is less reliable were excluded.

**Relationship of the** *C. bursa-pastoris* **subgenomes with their parental species.** To quantify the relationships between populations of *C. bursa-pastoris* and the two parental species, we applied a topology weighting method that calculates the contribution of each individual group topology to a full tree (34). We looked at the topologies joining each subgenome of *C. bursa-pastoris* and a corresponding parental lineage. There are 15 possible topologies for three populations of *C. bursa-pastoris*, a parental species, and the root. We grouped these topologies into five main groups: species trees - topologies that place a parental lineage as a basal branch to *C. bursa-pastoris;* three groups that join one of the populations of *C. bursa-pastoris* with a parental lineage and potentially signifies admixture; and all other trees that place a parental lineage within *C. bursa-pastoris* but do not relate it with a particular population of *C. bursa-pastoris* (Fig. 4A).

**Fig. 4.**
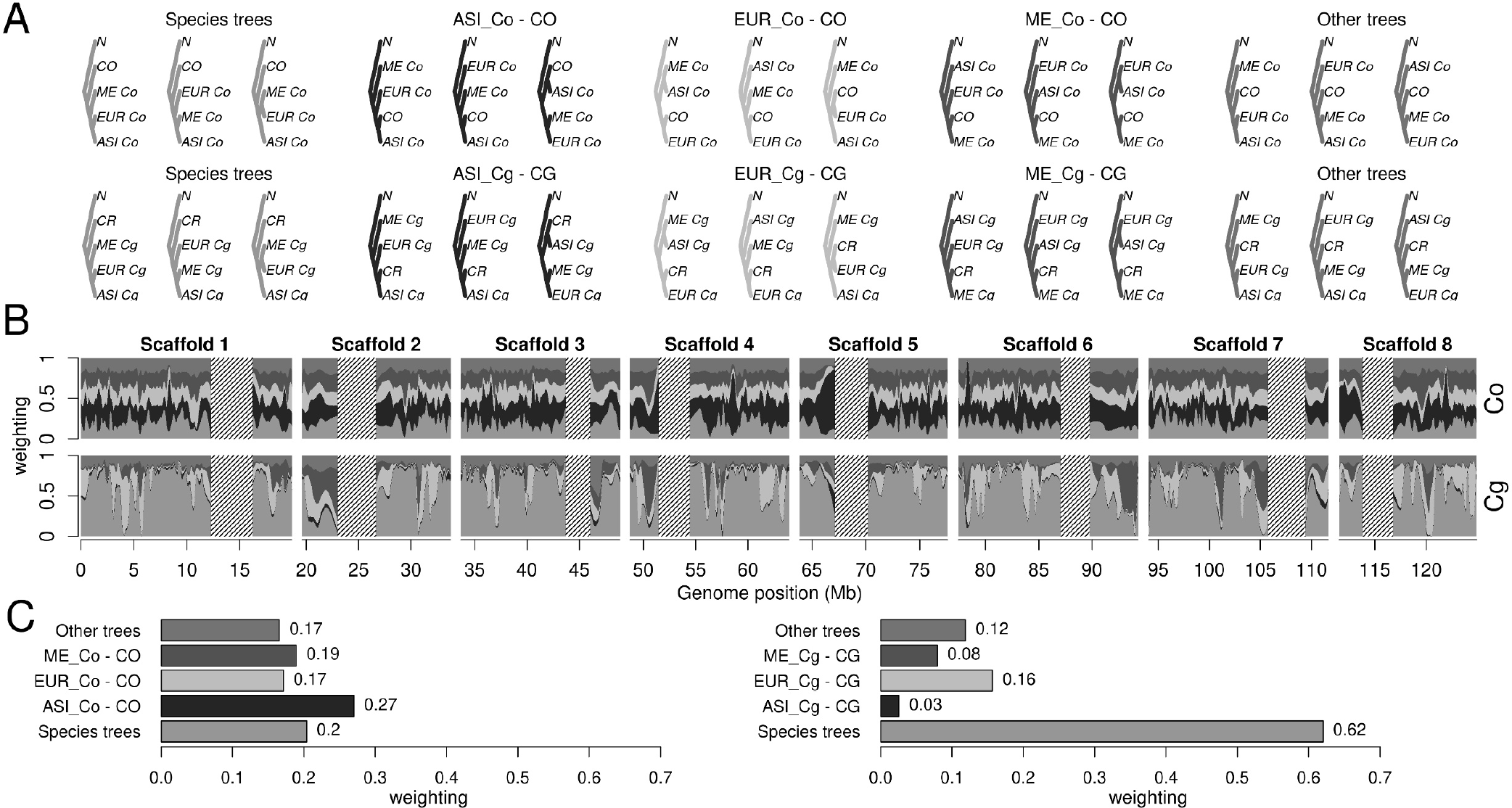
Topology weighting of the three populations of *C. bursa-pastoris* and parental species. (A) Fifteen possible rooted topologies for the three groups of *C. bursa-pastoris* in one subgenome and the corresponding parental species. The topologies are grouped into five main groups. Co and Cg indicate *C. orientalis* and *C. grandiflora /rubella* descendant subgenomes, respectively. ASI, EUR ME, CO, CR, N indicate Asian, European and Middle Eastern populations of *C. bursa-pastoris, C. orientalis, C. rubella*, and *N. paniculata*, respectively. (B) Topology weightings for 100 SNP windows plotted along 8 main scaffolds with loess smoothing (span = 1Mb). The tentative centromeric regions are shaded and only eight major scaffolds are shown. (C) Average weighting for the five main topology groups. The topology groups are in the same order (left-right and bottom-up) and colors in all plots.

These topology weightings varied along the subgenomes and illustrated distinct patterns between the two subgenomes (Fig. 4B). In the Co subgenome, the largest average weighting was for the topology grouping the ASI population of *C. bursa-pastoris* with *C. orientalis* (Fig. 4C), and the species topology had the second largest average weighting. The difference between the average weighting in these two topology groups was statistically significant (Table S3). In contrast, the species topologies weighting dominated in the Cg subgenome, regardless if *C. rubella* or *C. grandiflora* were used as a parental lineage (Fig. 4C, Fig. S3, Table S4, Table S5). The topology uniting the Cg subgenome of the EUR population with *C. rubella* was the largest among the topologies indicating admixture in the Cg subgenome (Fig. 4C). Thus, the two subgenomes differed substantially in the pattern of topology weighting and there were signs of a potential admixture of EUR and ASI with *C. rubella* and *C. orientalis*, respectively.

### Gene flow between *C. bursa-pastoris* and its relatives

#### Genomic inferences

The phylogenetic grouping of *C. orientalis* with the Asian Co subgenome, together with topology weighting results and the relatively elevated nucleotide diversity in this subgenome, suggested the presence of gene flow between *C. orientalis* and *C. bursa-pastoris* in the ASI population. To test this hypothesis, and at the same time to check for possibilities of gene exchange between *C. bursa-pastoris* and other *Capsella* species, we conducted two complementary tests of introgression.

We first used the ABBA-BABA test, a coalescent based method that relies on the assumption that alleles under incomplete lineage sorting are expected to be equally frequent in two descendant populations in the absence of introgression between any of them and a third population that diverged earlier on from the same common ancestor (35, 36). The deviation from equal frequency is measured with the *D-* statistics, which ranges between 0 and 1, with 0 indicating no gene flow and 1 meaning complete admixture. The ABBA-BABA test also provides an estimate of the fraction of the genome that is admixed by comparing the observed difference in ABBA-BABA with the difference expected under a scenario of complete admixture (*f*-statistics). We estimated *D* and f for triplets including one diploid species and two populations of *C. bursa-pastoris* represented by the most related subgenome to that species (Table 1). *N. paniculata* was the outgroup in all tests. The D-statistics were significantly different from 0 in most of the tests, so we considered all three combinations per test group (see Table 1) to determine the pairs that were the most likely to be admixed. The largest fraction of admixture was identified for the pair of the ASI Co subgenome and *C. orientalis* with an estimate of f indicating that at least 14% of the ASI Co subgenome is admixed. The second largest proportion of admixture was detected between *C. rubella* and the EUR Cg subgenome with f estimate of at least 8%. The estimates for tests with *C. grandiflora* were the smallest but similar to those obtained for *C. rubella*. The latter may reflect the strong genetic similarity between these two species rather than real gene flow between *C. grandiflora* and *C. bursa-pastoris* which, based on crosses (see below), seems unlikely. Finally, it should be pointed out that given that evidence for *C. bursa-pastoris* monophyly is weak, it is also possible that the signals of introgression from the parental species into *C. bursa-pastoris* that we are detecting here actually reflects introgression from an independently-arisen *C. bursa-pastoris* into either Co or Cg subgenomes.

We then used HAPMIX, a haplotype-based method, which should allow us to capture both large-scale and fine-scale admixture, and enables an absolute estimate of the proportion of the genome that was admixed. For the analysis of the Cg subgenome of *C. bursa-pastoris*, the highest levels of introgression were found consistently across regions to be from the diploid *C. rubella*. In Europe, 18% of SNPs genomewide showed introgression from *C. rubella*, followed by 11% in the Middle East, and just 2% in Asia (Table S6, Fig. S4A). All three populations also showed signs of *C. grandiflora* introgression but to a reduced extent compared to *C. rubella* (7% in Europe, 6% in the Middle East, 0.2% in Asia). *C. rubella* functionally represents a haplotype of *C. grandiflora*, and as noted above, we expect difficulties in discerning between the two, suggesting that much of the signal of introgression from *C. grandiflora* could in fact be due to *C. rubella* introgression. Of the regions putatively introgressed from *C. grandiflora*, 78%-96% of sites called as introgressed overlapped with those from *C. rubella*, none of which occurred in unique regions for *C. grandiflora*. Because of this, and in combination with the reduced genomewide probability of introgression from the diploid *C. grandiflora* compared to *C. rubella* (e.g. 0.11 compared to 0.24 in Europe), we argue that the signals of introgression from the diploid *C. grandiflora* were likely an artifact of its similarity with the regions of *C. rubella* introgression. These findings in accord with the ABBA-BABA results imply that the Cg subgenome has experienced significant introgression from *C. rubella* in Europe, and to a lesser extent in the Middle East.

For the analysis of the Co subgenome of *C. bursa-pastoris*, signals of introgression from the diploid *C. orientalis* were present in all three populations. In the ME population, 18-21% of SNPs showed signals of *C. orientalis* introgression (Table S6, Fig. S4B). Using the Middle East population for the analysis of the Co subgenomes of EUR and ASI, since it was the least introgressed in the HAPMIX results, yielded 15% *C. orientalis* introgression in Asia, and 14% in Europe. These findings suggest introgression of the diploid *C. orientalis* into the Co subgenome across all three geographic regions. Assuming these levels of admixture accurately reflect reality, we do not have a non-admixed reference population to use for Hapmix, and are thus violating a key assumption of the method. Hapmix inferences for the Co subgenome should therefore be taken with caution but we note that the results for ASI and ME are generally congruent with the admixture pattern obtained with ABBA-BABA.

#### Crosses

To assess further the plausibility of these inferences, we crossed individuals from the three populations of *C. bursa-pastoris* with their three diploid relatives to test for the presence of reproductive barriers. Regardless of the direction of the crosses, all crosses between *C. rubella* and the three populations of *C. bursa-pastoris* produced viable seeds. Importantly, crosses between *C. rubella* and EUR produced relatively more seeds and had smaller abortion rate than crosses with the other two populations of *C. bursa-pastoris*. Crosses between *C. orientalis* and *C. bursa-pastoris* mostly failed or led to aborted seeds, with the exception of one Russian accession of *C. orientalis* (PAR-RUS) that produced normally shaped seeds regardless if it served as a mother plant or as a pollen donor. In the latter case, there was a tendency towards higher seed number and smaller abortion rate for the ASI population than for EUR and ME. The crosses involving *C. grandiflora* mostly failed and the abortion rate approached 100%. Details on these crosses are provided in Appendix S1. Although the number of crosses was limited and did not provide enough power for proper statistical tests, they nonetheless were sufficient to show that the admixture detected at the molecular level was not completely restricted by reproductive barriers.

In summary, admixture between *C. bursa-pastoris* and *C. orientalis* in Asia, and between *C. bursa-pastoris* and *C. rubella* in Europe was supported by molecular data, even though some of the observed patterns could also be attributed to shared ancestry. Artificial crosses indicated that these inferences are credible.

### Selection and gene expression

#### Deleterious mutations

We first estimated the nucleotide diversity at 0-fold (*π*_0_) and 4-fold (*π*_4_) degenerate sites and then the ratio *π*_0_/*π*_4_ as a measure of purifying selection. Low values of *π*_0_/*π*_4_ would indicate higher purifying selection (37). As expected, *π*_0_/*π*_4_ was much lower in *C. grandiflora* than in *C. orientalis*. In *C. bursa-pastoris*, purifying selection was more efficient in the Cg subgenome than in the Co subgenome in both EUR and ME. However, the opposite was observed in the ASI population. For both subgenomes, the ASI population had the highest value of *π*_0_/*π*_4_ even if compared with *C. orientalis* (Fig. S5).

**Table 1.**
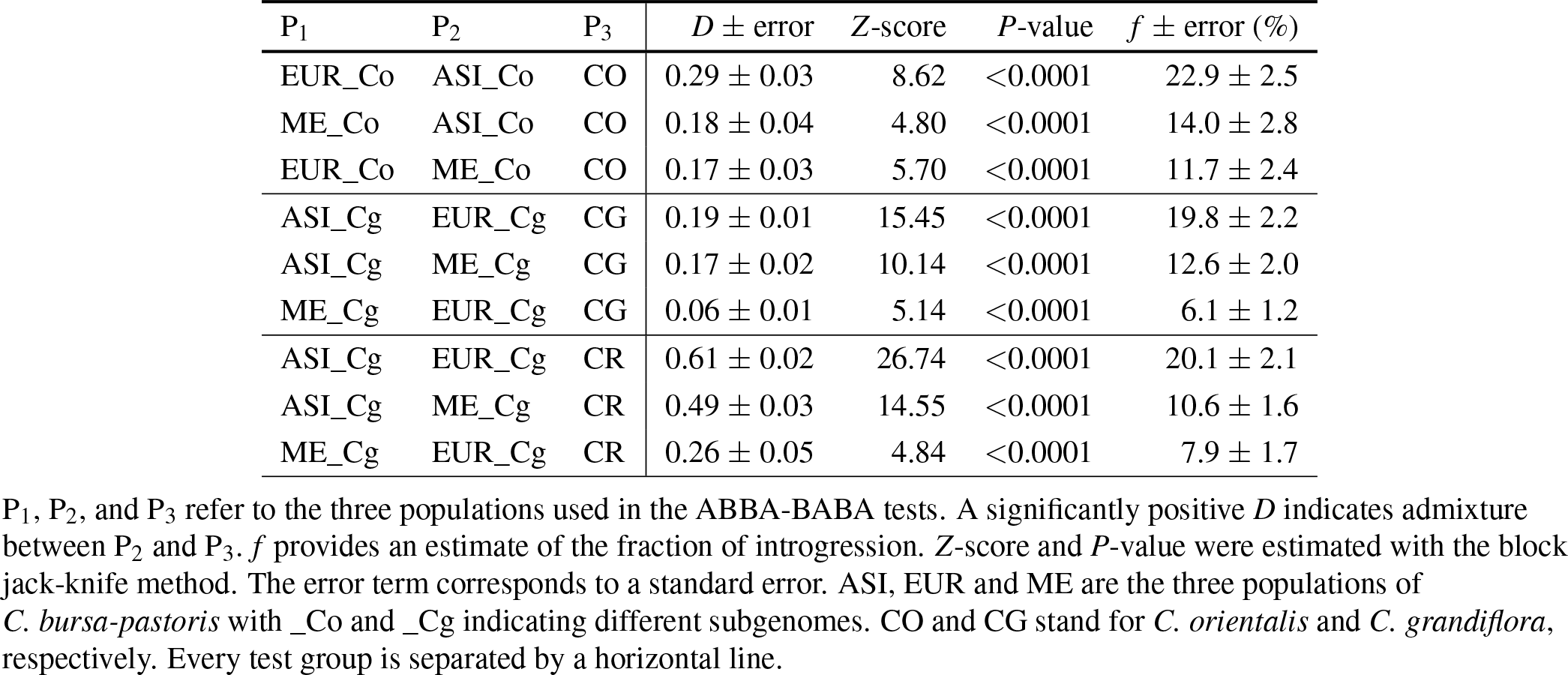
Results of the ABBA-BABA tests assessing admixture between *C. bursa-pastoris* and *C. orientalis, C. grandiflora* and *C. rubella*.

We then investigated the differences in deleterious mutations among subgenomes and populations by classifying nonsynonymous mutations with the SIFT4G algorithm that uses site conservation across species to predict the selective effect of nonsynonymous changes (38). In order to control for possible biases due to the unequal genetic distance between the different genomes and *C. rubella*, we used both *C. rubella* and *A. thaliana* SIFT4G annotation databases. Because we were interested in the number of deleterious mutations that accumulated after speciation of *C. bursa-pastoris*, we polarized the mutations of all three species with the reconstructed ancestral sequences of the common ancestors (see Material and Methods).

Regardless of the SIFT4G database (*C. rubella* or *A. thaliana*), the proportion of deleterious nonsynonymous sites among derived mutations was always significantly higher in *C. orientalis* and the Co subgenomes than in *C. grandiflora* and the Cg subgenomes (Fig. 5, Table S7, Table S8). Within *C. bursa-pastoris*, the proportion of deleterious mutations depended on the population considered with the highest value in the ASI population and the smallest in EUR. It is also noteworthy that the proportion of deleterious nonsynonymous sites of the Co subgenome in EUR and ME was significantly smaller than that of *C. orientalis* suggesting that a higher effective population size in the Co subgenome than in its ancestor led to more efficient purifying selection in these two populations. On the other hand, the proportion of deleterious nonsynonymous sites in the Asian Co subgenome was larger than in *C. orientalis*, but this difference was only significant for the *A. thaliana* database. The Cg subgenome also had a significantly higher proportion of deleterious sites in ASI than in EUR and ME in all comparisons. In conclusion, the proportion of deleterious sites in the two subgenomes of extant *C. bursa-pastoris* still reflected the differences between the parental species and the efficacy of purifying selection in the different *C. bursa-pastoris* populations was associated to their synonymous nucleotide diversity or, equivalently, to their effective population size.

**Fig. 5.**
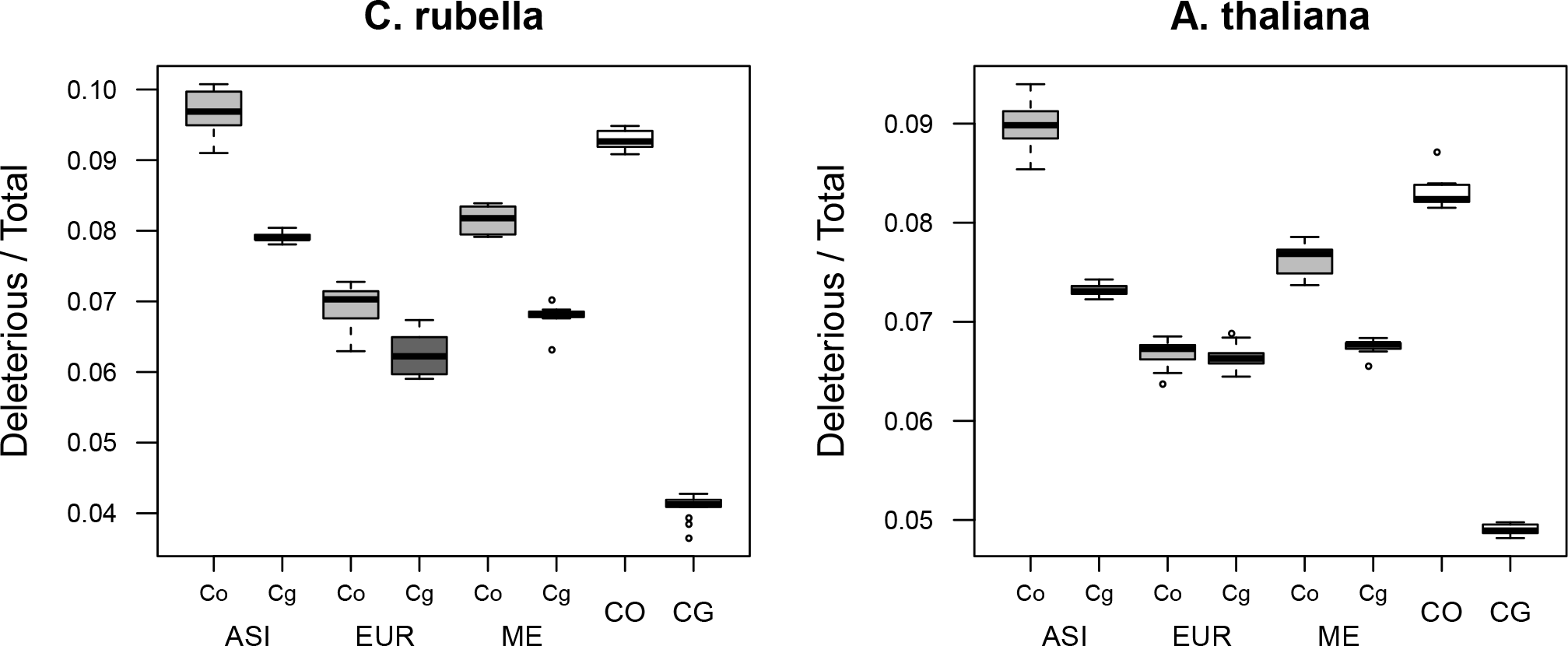
Genetic load in the subgenomes of *C. bursa-pastoris* and its parental species. The proportion of deleterious nonsynonymous changes was estimated with SIFT4G on derived alleles, i.e. alleles accumulated after the speciation of *C. bursa-pastoris*. The left plot shows the results obtained with *C. rubella* database and the right plot those obtained with *A. thaliana* database. Co and Cg are the two subgenomes of *C. bursa-pastoris*. ASI, EUR, ME, CO, CG indicate Asian, European and Middle Eastern populations of *C. bursa-pastoris*, and parental species *C. orientalis* and *C. grandiflora*, respectively.

#### Selective sweeps

The three populations of *C. bursa-pastoris* also differ in patterns of positive selection. Overall, the number of sweeps in Co and Cg subgenomes were independent (*χ*^2^ = 89.386, p-value < 0.001). Selective sweeps were more significant on the Cg subgenome than on the Co subgenome in EUR and ME, whereas in the ASI population, the opposite was true (Fig. 6). The regions harboring significant sweeps were also larger on the Cg subgenome than on the Co subgenome in EUR and ME (total length 42 Mb, 50 Mb vs 9 Mb, 3 Mb), whereas in Asia the sweep regions were larger on Co than on Cg (total length 4 Mb vs 830 Kb). Although the locations of the Cg sweeps in EUR and ME largely overlap, the patterns differed between the two populations. For example, the strongest sweep in EUR was located on scaffold 1, whereas the strongest sweep in ME was on scaffold 6. In addition, EUR had many sweeps in Co subgenome (109 in EUR_Cg, 128 in EUR_Co), but they all were small and hardly above the significance threshold (Fig. 6). In the ME population, the sweeps in the Cg subgenome were prevailing both in size and numbers (101 in ME_Cg, 22 in ME_Co). The ASI population differed strongly from both EUR and ME not only because most of its sweep signals were on the Co subgenome but also because these sweeps regions were narrower and less pronounced (Fig. 6). Thus, all three populations of *C. bursa-pastoris* were distinct in their selective sweeps patterns with the Asian population being the one least affected by positive selection.

**Figure 6.**
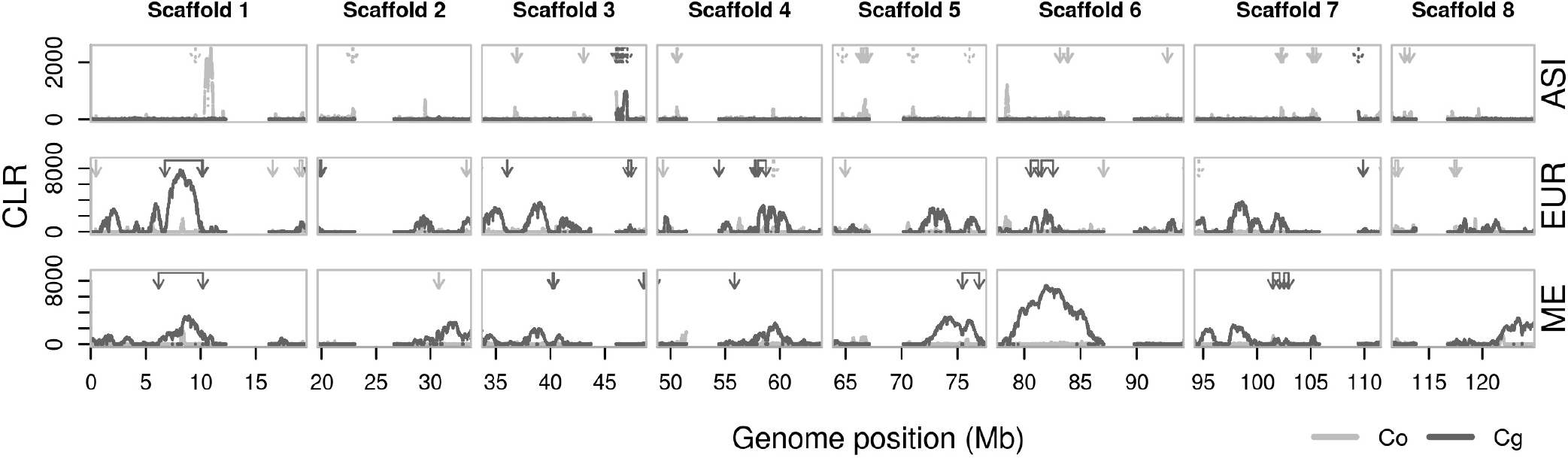
Selective sweep differences between populations of *C. bursa-pastoris*. Selective sweeps are detected with the composite likelihood ratio statistics (CLR) along the Co and Cg genomic subgenomes in Asian (ASI), European (EUR) and Middle Eastern (ME) populations. Solid arrows point to the location of introgression and dashed arrows show the location of genomic conversion. Pericentromeric regions are removed. Only eight major scaffolds are shown.

Given the presence of gene flow between *Capsella* species, we also checked if any of the detected selective sweeps could be due to introgression. We compared genetic distances for every sweep region among the three *Capsella* groups and the parental species. A sweep region was considered to have resulted from introgression if its genetic distance was closer to the corresponding parental species than to any other *C. bursapastoris* sequence. This comparison also allowed us to identify regions of possible gene conversion if the genetic distance was the smallest between the two subgenomes. The distance between individual sweep regions revealed that all the composite likelihood ratio (CLR) outliers signifying sweeps in the ASI Cg subgenome were genetically closer to the Asian Co subgenome than to other Cg subgenomes of *C. bursapastoris* (Fig. S6), and thus they probably were the result of gene conversion. The distance analysis of sweep regions in the ASI Co subgenome revealed 9 regions of gene conversion (total length 505 Kb) and 17 regions of introgression from *C. orientalis* (total length 1.3 Mb) (Fig. 6, Fig. S6). On the other hand, we found 9 regions of potential introgression from *C. orientalis* to the EUR Co subgenome (total length 945 Kb), and one to the ME Co subgenome (length 40 Kb). There were also 10 introgressions between *C. rubella* and the EUR Cg subgenome (total length 6.5 Mb), and 7 introgressions between *C. rubella* and the ME Cg subgenome (total length 6.7 Mb) (Fig. S6). We did not observe any sign of gene conversion in the ME population and in the EUR Cg subgenome, but we found 2 regions of gene conversion from the Cg to the Co subgenome in EUR (total length 154 Kb). The regions of gene conversion showed reduced heterozygosity in both the phased and unphased data (Fig. S7), suggesting they were not an artifact of phasing. Thus, some of the sweep signals could be solely due to gene conversion and introgression, but we cannot rule out subsequent selection of these conversion and introgression regions.

#### Homeologue-specific expression

The relative expression of the two subgenomes, or homeologue-specific expression (HSE), can provide additional information on the evolution of the two subgenomes in different populations of *C. bursa-pastoris*. In particular, biased adaptation towards one subgenome may select for decreased expression of the other subgenome. Given selective favor for different subgenomes in different populations, one would also expect the Cg subgenome to be over-expressed in EUR and ME, and the Co subgenome in Asia.

To assess HSE, we analyzed the RNA-Seq data of 24 accessions representing all three populations of *C. bursa-pastoris* in a hierarchical Bayesian model that integrates information from both RNA and DNA data (39). Overall, in agreement with Douglas et al. (9), one subgenome did not dominate the other in the 24 accessions considered together, though a few genes demonstrated a slight expression shift toward the Cg subgenome. On average, we assessed HSE in 13,589 genes per accession (range 12,808-15,340) and 18% of them showed significant HSE (posterior probability of HSE > 0.99). The expression ratios between subgenomes (defined here as Co / Total) across all assayed genes in the DNA data were close to equal (mean = 0.495). Thus, there was no strong mapping bias.

Among populations, HSE varied considerably. The mean expression ratios for all genes were 0.494, 0.489, and 0.489 in the ASI, EUR, and ME accessions, respectively, and these mean ratios for genes with significant HSE were 0.487, 0.465, 0.468. The difference in mean ratio between EUR and ME was not significant, but both EUR and ME were significantly different from ASI (Table S9). In addition, the distribution of expression ratios between the two subgenomes was right-skewed in EUR and ME, whereas in the ASI population, the distribution was more symmetrical (Fig. 7). The difference between the populations was particularly evident in the grand mean values (Fig. 7). Thus, the shift towards higher expression of the Cg subgenome was more prominent in Europe and the Middle East than in Asia.

**Fig. 7.**
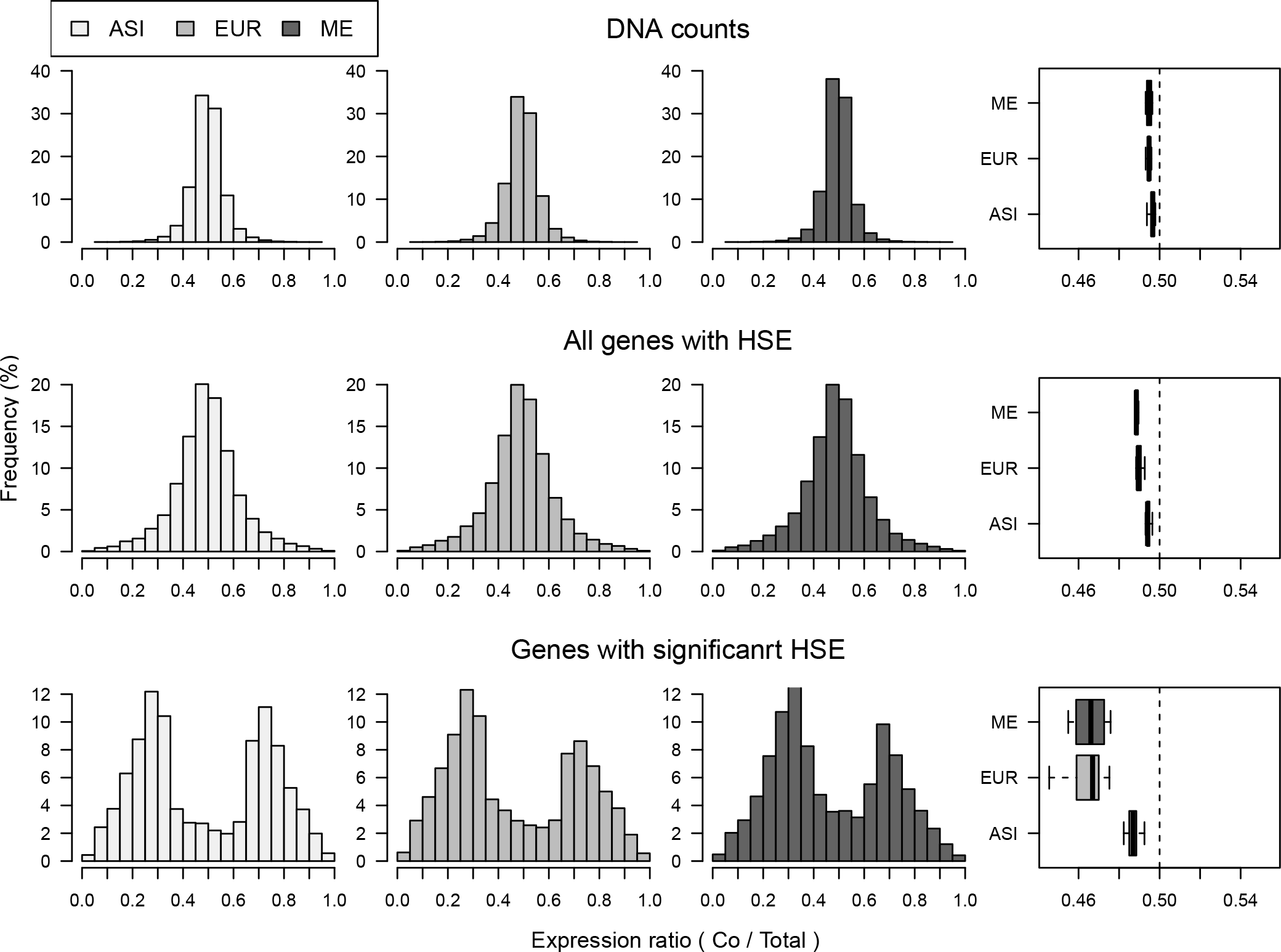
Distributions of expression ratios between the two subgenomes of *C. bursa-pastoris*. The subgenome specific expression (HSE) is estimated by the fraction the Co subgenome relative to total expression level. The upper part presents the distributions for DNA counts, the middle plots show the expression distribution for all assayed gene and the lower plot shows only the distribution for genes with significant expression of one of the subgenomes. The histograms present the distribution of allelic ratio, whereas the boxplots summarize these results with the grand mean for every sample. ASI, EUR, and ME indicate Asian, European and Middle Eastern populations, respectively.

Expression levels were also noticeably distinct in the three populations. We analyzed the pairwise correlations in the HSE between all 24 samples, to check if the direction of the expression shift in every gene was similar within and between populations. Overall, the levels of expression of the genes with significant HSE were positively correlated between samples (mean Pearson’s *r*=0.81), but correlations were distinctly stronger for samples from the same population (mean Pearson’s *r* = 0.91) than for samples from different populations (mean Pearson’s *r* = 0.75) (Fig. S8). This pattern was also similar for the pairwise correlations across all assayed genes but the correlation coefficients were smaller (mean Pearson’s *r* for all comparisons 0.56, within populations 0.72 and between populations 0.47) (Fig. S9). Thus, globally expression levels co-varied, but they were more similar within populations than between them.

The genes with significant HSE were roughly the same in all three populations. We considered that a gene showed a population-specific HSE if it had a significant HSE in at least 9/11, 7/9, and 2/4 samples for ASI, EUR, and ME, respectively. With these criteria, we found that there was almost 60% overlap in gene names showing significant HSE in pairwise comparisons between the three populations. Also, selective sweep regions were not over-represented by genes with significant HSE (Fisher’s Exact Test, p-values 0.99, 0.91, 0.47 for ASI, EUR, ME, respectively.).

Additionally, we were interested in testing whether the results of the differential gene expression analysis of phased data between these three populations differed from the results obtained by Kryvokhyzha et al. (32) on unphased data. Many genes differentiated the ASI, EUR and ME populations in Kryvokhyzha et al. (32), but all differences could be explained by population structure. We performed similar tests on the phased data and obtained almost the same results (see Appendix S2). The ASI and EUR populations showed the largest number of genes differentially expressed, and EUR and ME the smallest. However, this pattern was not detectable in the model accounting for population genetic structure. Thus, variation in expression level based on phased data between two subgenomes did not differ much from the variation based on unphased data and could as well be explained by the demographic processes in these populations.

## Discussion

In the present study, we analyzed the genetic changes experienced by a recently formed allopolyploid *C. bursa-pastoris* since its founding, focusing on the evolutionary trajectories followed by its two subgenomes in demographically and genetically distinct populations from Europe, the Middle East, and Asia. The shift to selfing and polyploidy had a global impact on the species, resulting in a sharp reduction of the effective population size in all populations, that was accompanied by relaxed selection and accumulation of deleterious mutations. However, the two subgenomes were not similarly affected, with the magnitude of the subgenome-specific differences depending on the population considered. The relative patterns of nucleotide diversity, genetic load, selection and gene expression between the two subgenomes in the European and the Middle Eastern populations were distinct to that observed in Asia. The differences between populations were further enhanced by post-speciation hybridization of *C. bursa-pastoris* with local parental lineages. Below, we discuss these global and local effects in more detail and their consequences for the history of the species.

### Effect of parental legacy

The effective population size of the diploid outcrossing ancestor of *C. bursa-pastoris, C. grandiflora*, is ten times larger than that of its selfing ancestor *C. orientalis* (9). Any analysis of the difference in effective population size between the subgenomes of *C. bursapastoris* or of their evolutionary trajectories must therefore account for this initial difference. After the bottleneck associated with the origin of *C. bursa-pastoris* and the reduction in *N_e_* due to the shift to selfing (40), the effective population sizes of the two subgenomes are expected to progressively converge and decrease along the same trajectory.

While this was indeed the observed overall pattern, the trajectories followed by the two subgenomes in the three populations differed: in Europe the initial level was similar to that in the Middle East but higher than in Asia and the decline of *N_e_* of the Cg subgenome was delayed compared to the sudden decline experienced by the Co subgenome. In contrast, in Asia the two subgenomes initially followed similar downwards trajectories but *N_e_* increased again in both subgenomes at around 40,000 ya. In the diploid *C. orientalis*, there was a period of stasis followed by a steeper decline than in the tetraploid. The difference in demography across the three regions could indicate multiple origins of *C. bursa-pastoris* as suggested by Douglas et al. (9) and the difference between the diploid and the tetraploid could reflect a mixture of the population expansion experienced by the tetraploid and the buffering effect of tetraploidy against deleterious mutations.

There was a clearly noticeable difference between the two subgenomes in the number of inherited deleterious mutations. Based on the strong differences in *N_e_*, one would expect the efficacy of selection to be much higher in *C. grandiflora* than in *C. orientalis* that has a much smaller *N_e_*, (41). In the analysis of the genetic load, we indeed observed that *C. orientalis* had a higher proportion of deleterious mutations than *C. grandiflora*. Hence, the amount of genetic load most likely was different between the Cg and Co subgenomes of *C. bursa-pastoris* at the time of the species emergence. Interestingly, hundreds of thousands of generations of selfing did not totally erase the differences between the two subgenomes and, today, the Co subgenome still carries more deleterious mutations than the Cg subgenome. This difference was smaller than the difference between *C. orientalis* and *C. grandiflora*, but it was still significant. Nucleotide diversity also demonstrated the effect of parental legacy. The Cg subgenome inherited from the more variable outcrosser *C. grandiflora* was still more diverse in all populations except the Asian one. The maintenance of part of the parental legacy in both cases suggest that, in spite of their initial differences, both subgenomes have experienced similar levels of fixation since the creation of the species. The Asian population is an exception in this regards because it was affected by secondary gene flow as discussed below. Variation in nucleotide diversity in the coding part of the genome also demonstrated similarity in the efficacy of purifying selection between the two subgenomes and their corresponding parental lineages, though the pattern in the ASI population was the reverse of that observed in the parental lineages. The effect of parental legacy on gene degeneration was also noted in Douglas et al. (9). Thus, the effect of the genetic background of hybridizing species may not be as overwhelming as the effect of mating system but it still impacts the fate of the two subgenomes long after the species arose.

### Subgenome-specific introgression and/or multiple origins

Based on coalescent simulations and the amount of shared variation between *C. bursa-pastoris* and its parental species Douglas et al. (9) ruled out a single founder but noted that it would be very difficult to estimate the exact number of founding lineages. Douglas et al. (9) did not consider hybridization but an earlier study detected gene flow from *C. rubella* to the European *C. bursa-pastoris* using 12 nuclear loci and a coalescent-based isolation-with-migration model (42). The present study adds two new twists to the story. First, our results indicate that shared polymorphisms were not symmetrical: namely, in the EUR and ME populations introgression from *C. rubella* occurred on the *C. grandiflora* subgenome whereas in ASI introgression from *C. orientalis* occurred on the *C. orientalis* subgenome. Second, in both the whole genome and density trees, *C. orientalis* appears as derived from the *C. bursa-pastoris* Co subgenome rather than the converse as one would have expected. No such anomaly was observed for *C. grandiflora* that, as expected, grouped at the root of the *C. bursa-pastoris* Cg subgenome. These results could be explained by a mixture of multiple origins and more recent introgression. Multiple origins seem to be common in allotetraploids (23, 43) and interploidy gene flow has already been inferred for the *Capsella* (42) and other plant genera (44, 45).

Our crossing results did not reject the possibility of ongoing admixture between *C. bursa-pastoris* and parental lineages in both Europe and Asia. European and Asian populations of *C. bursa-pastoris* partially overlap in the distribution ranges with *C. rubella* and *C. orientalis*, respectively (Fig. 1A). The exact proportion of introgression remains unclear at this stage. Taken at face value, the strongest admixture was between the ASI Co subgenome and *C. orientalis*. Considering the overlapping estimates of *f*-statistics and HAPMIX, the proportion of admixture of the ASI Co subgenomes with *C. orientalis* was around 14%-23%. The admixture between the EUR Cg subgenome and *C. rubella* was also strong, being around 8-20%. There were also signs of minor admixture in the ME population with both *C. orientalis* and *C. rubella*. This lack of a non-admixed population posed a problem of correct estimation of the proportion of admixture for both the ABBA-BABA and HAPMIX approaches.

In the ABBA-BABA test, departure from the assumptions can lead to under-or overestimated introgression. In the present case, some proportion of the variation shared between *P*_3_ and both *P*_1_ and *P*_2_ populations could be due to introgression and not to incomplete lineage sorting and this would lead to underestimating the amount of admixture. On the other hand, small *N_e_* and recent divergence of the populations used in the test can inflate estimates of *D* (46). Further, the behavior of *D* in tests involving both selfing and outcrossing species has not been assessed yet. The *D* statistics were significantly different from zero in all our comparisons suggesting that admixture did indeed occur in all populations of *C. bursa-pastoris*. The f statistic is considered less prone to be affected by these factors (46), and it was more reliable in our tests too. Its values were close to zero in the alternative combinations for the ABBA-BABA tests where we did not expect to find admixture, while *D* had high estimates (Table S10). Thus, the f values are the closest to the real proportion of admixture we could get.

In HAPMIX, when one reference population is admixed, the program probably compensates for this extra relatedness between the reference populations by inflating intermediate introgression probabilities. Therefore, we observed the discrepancy between the results of HAPMIX and ABBA-BABA in the estimates of admixture between the EUR Co subgenome and *C. orientalis*. However, the results for the Cg subgenome largely agreed between HAPMIX and ABBA-BABA and, together with the results by Slotte et al. (42) and our crossing experiment, bolsters the hypothesis of admixture between *C. rubella* and *C. bursa-pastoris* in Europe. On balance, a scenario with a single origin of *C. bursa-pastoris* with later rampant admixture with *C. orientalis* in Asia and less extensive admixture with *C. rubella* in Europe is consistent with our data.

On the other hand, our results could also be obtained under a scenario of multiple origins. This scenario seems particularly likely if one looks at Fig. 4, where the histories of the Co and Cg subgenomes are totally different. If we assume that *C. orientalis* and *C. grandiflora* are indeed parental lineages and there was no unknown parental lineage that went extinct, this picture can be only explained by a separate and more recent origin of the ASI population (Fig. 8). However, the scenario of multiple origins and post-speciation admixture are not mutually exclusive. The signs of gene flow between EUR and *C. rubella* are still best explained by post-speciation admixture. The weak signs of admixture between *C. bursa-pastoris* and *C. orientalis* in EUR and ME are also difficult to fit into a scenario involving only multiple origins. A possibility is that these signs of admixture resulted from gene flow from ASI to EUR and ME within *C. bursa-pastoris*. The ASI population is more related to *C. orientalis* and the presence of its alleles in EUR and ME could be spuriously recognized as introgressed from ASI. Regardless of whether a single or a multiple origin scenario is the true one, our results demonstrate that the history of *C. bursa-pastoris* is far more complex than previously imagined.

**Fig. 8.**
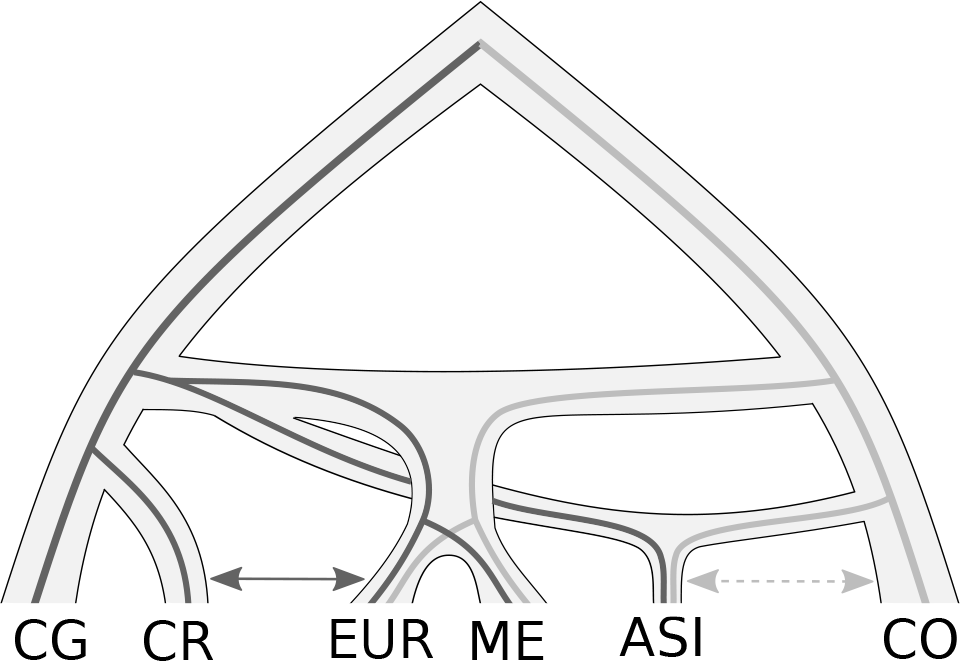
A tentative scenario of multiple origin of *C, bursa-pastoris*. The Asian population originated separately from other *C, bursa-pastoris* populations. There may still be gene flow between the Asian population and *C, orientalis* (dashed arrow). There is gene flow between the European *C, bursa-pastoris* and *C, rubella* (solid arrow). ASI, EUR, ME, CR, CO, CG indicate Asian, European and Middle Eastern populations of *C, bursa-pastoris, C, rubella*, and parental species *C, orientalis* and *C, grandiflora*, respectively.

### Weak subgenome-specific expression differences

Many allopolyploid species show subgenome expression bias, where one subgenome tends to be over-expressed relative to the other one (47–50). This expression dominance is often observed in synthetic allopolyploids (51–54) and thus the major part of such preferential subgenome dominance is probably established immediately after allopolyploidization. The subgenome expression dominance is also suggested to be largely defined by parental expression differences (55, 56). Contradictory results on patterns of subgenome specific expression in *C. bursa-pastoris* have been obtained so far. Douglas et al. (9) concluded that there is no strong homeologue expression bias and those few genes showing HSE could be explained by parental expression differences. However, genes with HSE do show a slight bias towards overexpression of the Cg subgenome inherited from *C. grandiflora / rubella* lineage on the Figure 3B in Douglas et al. (9). In contrast, Steige et al. (57) reported a higher expression of the Co subgenome inherited from *C. orientalis* in three accessions, and Cg over-expression in a fourth one (CbpGR). Steige et al. (57) hypothesized that the over-expression of the Co subgenome might be related to a higher number of transposable elements in this subgenome, but they did not find any evidence of this and could not explain the down-regulation of the Co subgenome in the CbpGR accession and in the artificial hybrid between *C. rubella* and *C. orientalis*.

Considering the population histories of *C. bursa-pastoris* sheds some light on these discrepancies. The results of Douglas et al. (9) and Steige et al. (57) are consistent with the hypothesis that *cis*-regulatory differences between the *C. orientalis* and *C. grandiflora / rubella* genomes result in overexpression of the Cg subgenome in a hybrid comprising both genomes. Thus, in the absence of other factors, the slight over-expression of the Cg subgenome would be the default HSE pattern in *C. bursa-pastoris*. In accordance with this, we observed over-expression of the Cg subgenome in the ME and EUR populations that are most likely the closest to the region of origin of *C. bursa-pastoris* (31). The accessions that show over-expression of the Cg subgenome in Douglas et al. (9) (SE14 from Sweden) and in Steige et al. (57) (CbpGR from Greece), as we now know belong to the EUR population (31). Hence, their results are consistent with ours and expected if the HSE is defined primarily by the differences between the parental lineages. On the other hand, we observed that genes with HSE in the ASI population showed equal expression between the two subgenomes. The accessions showing over-expression of the Co subgenome in Steige et al. (57) also mostly belong to the ASI population (CbpKMB and CbpGY, though not CbpDE that putatively originates from Germany). Thus, the Asian accessions show the HSE that is different from the default pattern. This difference can be caused by the selection preference for the Co subgenome and/or by introgression from *C. orientalis* that enhanced the *cis*-regulatory elements of the Co subgenome. The ASI population experienced a strong population bottleneck, so genetic drift played some role as well. These explanations need to be confirmed because HSE can be influenced by many factors (e.g. *trans*-regulatory elements, gene methylation, transposable elements), but it is clear that there are different directions of HSE in populations of *C. bursa-pastoris* and they are caused by the different evolutionary histories of those populations.

The reason we observed an equal expression between subgenomes in ASI, whereas Steige et al. (57) detected expression bias of the Co subgenome for Asian samples, could also be due to different approaches in our analyses. First, we extracted RNA from seedling, whereas Steige et al. (57) obtained RNA from leaves and flower buds. Variation in HSE for different tissues of *C. bursa-pastoris* is not characterized yet, so the Co expression in seedlings may not be apparent yet. Second, we mapped reads to the *C. rubella* reference with masked polymorphism, whereas Steige et al. (57) used the reconstructed reference of an F1 hybrid between *C. orientalis* and *C. rubella*. The bias in our DNA data was not stronger than in Steige et al. (57), so which method is more appropriate remains to be found out.

### Neutral inter-population expression differences

We have previously reported that differences among populations in overall gene expression variation (i.e. from unphased data) in *C. bursa-pastoris* primarily reflect population structure and hence are mostly driven by genetic drift (32). The current study of phased gene expression data is consistent with this result. Both the differential gene expression analysis of each subgenome and the generalized linear model analysis of HSE data as proportions revealed similar differences between populations and these differences were all explained by the genetic population structure in the species. Our results also demonstrated that genes showing significant HSE largely overlapped between populations and these genes were not strongly enriched for GO terms. These genes probably evolve under a compensatory drift model (58). This was evident in the direction of the HSE, which was the same in all accessions. The correlation in levels of HSE is stronger within than between populations, which is also consistent with evolution by drift. Hence, gene expression variation does not show strong adaptive changes in the early stages of the evolution of *C. bursa-pastoris*. It is still possible that some of the gene expression differences are not neutral and we have previously discussed the potential pitfalls of detecting adaptive differences in structured populations (32). The asymmetric over-expression between populations, for instance, agrees with the presence of some selective differences between populations.

## Conclusion

Three salient, and sometimes unexpected, features of the evolution of the tetraploid shepherd’s purse that emerged from the present study, are its complex origin and the magnitude of introgression with diploid relatives, the long-lasting effects of the difference between its two parental species and the importance of demography in shaping its current genomic diversity. Hence, the present study suggests that understanding the evolution of tetraploid species without paying due attention to the historical and ecological backgrounds under which it occurred could be misleading.

## Materials and methods

### Sequence data

We obtained the whole genome sequences of 31 accessions of *C. bursa-pastoris* and the seedling transcriptomes of 24 of these accessions. Transcriptome data used in this study were generated previously (32). Whole genome DNA data consisted of 10 accessions downloaded from GenBank (PRJNA268827) and 21 accessions sequenced in this study. New DNA samples were sequenced using the same technology as the downloaded ones (100- bp paired-end reads, Illumina HiSeq 2000 platform, SciLife, Stockholm, Sweden). The mean genomic coverage of *C. bursa-pastoris* samples was 47x. We also used genomic data of 10 *C. orientalis* and 13 *C. grandiflora* samples from GenBank (PRJNA245911, PRJNA254516). For the analysis requiring an out-group, we used the whole genome assembly of *Neslia paniculata* (59). Detailed information on the samples is provided in Table S1.

### Genotype calling and phasing

DNA reads from each individual were mapped to the *Capsella rubella* reference genome (59) using Stampy v1.0.22 (60) with default parameters, except that the substitution rate was set to 0.025 to account for the divergence from the reference. Potential PCR duplicates were marked using Picard Tools 1.115 (http://picard.sourceforge.net) and were ignored during genotyping. Genotypes were called using *HaplotypeCaller* from the Genome Analysis Tool Kit (GATK) v3.5 in the GVCF mode and heterozygosity set to 0.015 (61). Genotypes were filtered for depth between 6 and 100 reads (the 5th and 99th coverage percentiles, respectively). This approach produced a VCF file containing all called sites. This VCF was used in the analyses requiring both polymorphic and monomorphic sites for correct estimates. To obtain a set of SNPs with the highest confidence possible, we generated another VCF file that contained only polymorphic sites and applied more stringent filtering. We set to no-call all sites that met the following criteria: *MQ < 30, SOR > 4, QD < 2, FS > 60, MQRankSum < −20, ReadPosRankSum < −10, ReadPosRankSum > 10*. These filtering criteria were defined following GATK Best Practices (62) with some adjustment guided by the obtained distributions of the GATK annotation scores (Fig. S10).

To phase the *C. bursa-pastoris* subgenomes, we run HapCUT version 0.7 (63) on each sample from the VCF with the stringently filtered SNPs. The phased haplotype fragments were then joined into two sequences descended from *C. grandiflora* and *C. orientalis*. The origin of haplotypes in HapCUT fragments was defined using sites with fixed heterozygotes in *C. bursa-pastoris* and fixed differences between *C. grandiflora* and *C. orientalis*. Fragments that had small (< 2 sites) or no overlap with variation in *C. grandiflora* and *C. orientalis* as well as those that looked chimeric (prevailing phasing state was supported by less than 90% of sites) were set to missing data (Fig. S11). Additionally, we also set to missing the sites that were defined as not real variants or not heterozygous by HapCUT (flagged with *FV*). HapCUT phasing produced the alignment that had only heterozygous sites and removed all the sites that were non-variant within but variable between individuals. We restored this inter-individual variation with introduction of the same proportion of missing data into non-variant sites as it was introduced to heterozygous sites during the phasing. Similarly, we also merged the phased SNPs dataset with whole genome data.

The reference genomes of *C. grandiflora* and *C. orientalis* were created using the GVCF files produced by Douglas et al. (9). The variants were called as described above with additional filtering for fixed differences between the two species. For some of the analyses, where the software was not able to treat heterozygous genotypes properly, we pseudo-phased the sequences of *C. grandiflora* and *C. orientalis* by randomizing alleles in heterozygous genotypes.

The final data-sets in all the analyses comprised the alignment of phased *C. bursa-pastoris* sequences, *C. grandiflora, C. orientalis, C. rubella* (the reference sequence) and *N. paniculata*. This alignment was filtered for missing data such that genomic positions with more than 80% of missing genotypes were removed. We also removed the repetitive sequences as annotated in Slotte et al. (59) and pericentromeric regions that we delineated based on the density of repetitive regions and missing data.

### Reconstruction of the ancestral sequences

Several analyses presented in this paper required polarized sequence data. The most common approach to polarizing the alleles is to use an outgroup. However, the alignment of *Capsella* species and *N. paniculata*, the nearest outgroup with a whole genome sequence available, resulted in substantial reduction of the dataset due to missing data. To overcome this drawback, as well as to track mutations’ origin on the phylogenetic branches, we reconstructed ancestral sequences for major phylogenetic splits. The reconstruction was performed on the tree that was assumed to represent a true history of the *Capsella* species (Fig. S12) using the empirical Bayes joint reconstruction method implemented in PAML v4.6 (64).

### Population differentiation

To assess the degree of differentiation among populations for the two subgenomes, we estimated absolute divergence (*Dxy*) and nucleotide diversity (7r) of the phased genomes using a sliding window approach. The estimates were calculated on non-overlapping 100 Kb windows using the *EggLib* Python module (65). The *p-* values for the difference in mean values were estimated using 10,000 bootstrap resamples from 100 Kb windows.

### Temporal change in *N_e_*

We reconstructed changes of *N_e_* over time with both PSMC (66) and SMC++ (33). We first masked potential CpG islands and all nonsynonymous sites in the genome to avoid bias caused by variation in mutation rates or selective effects. We randomly paired haplotypes for estimation in *C. orientalis* and did the same for estimations based on the two subgenomes of *C. bursa-pastoris*. SMC++ was run on all samples from a population, with default parameter settings. For PSMC runs, we set parameters to “-N25-t15-r5-p 4+25*2+4+6”. Variation in *N_e_* was estimated using 100 bootstrap replicates and three different pairs. We chose a mutation rate equal to the mutation rate of *A. thaliana*, *μ* = 7 × 10^-9^ per site per generation (67) and a generation time of 1 year for all *Capsella* species.

### Phylogenomic analyses

We reconstructed a whole genome phylogeny to explore the relationship between the phased subgenomes of the three populations of *C. bursapastoris* as well as its parental species. To investigate the local phylogenetic relationships along the genome, we also conducted a sliding window phylogenetic analysis using non-overlapping 100 Kb windows. In both analyses, phylogenetic trees were reconstructed using the neighbor-joining algorithm and absolute genetic distance in R package ape (68). Additionally, a whole genome phylogenetic tree was also reconstructed using the maximum-likelihood approach with the GTRGAMMA model and 100 boostrap replicates in *RAxML* v8.2.4 (69) (Fig. S13). The trees from the sliding window analysis were described by counting the frequency of monophyly of different groups with the Newick Utilities (70). The variation in topology across the genome was also described using topology weighting implemented in TWISS (34). The weighting was estimated for 100 SNPs windows where each sample was genotyped for at least 50 SNPs. To test for the difference in mean topology weighting, we fitted the generalized linear model with a binomial distribution and performed multiple comparisons for the contrasts of interest with the *glht* function from the *multcomp* library in *R* (71).

### Tests for gene flow

To evaluate the presence of gene flow between the parental species and *C. bursa-pastoris*, we calculated the ABBA-BABA based statistics, D, an estimate of departure from incomplete lineage sorting, and f, an estimate of admixture proportion (35, 36). These statistics and their significance, which was estimated with a 1Mb block jackknife method, were calculated from population allele frequencies with scripts from Martin et al.(72). We also used HAP-MIX (73) to infer haplotype blocks of introgression from the diploids *C. grandiflora, C. rubella*, and *C. orientalis* into the three populations of *C. bursa-pastoris* for each phased subgenome. We removed sites with more than 20% missing data for each population. The remaining missing data was imputed for the parental populations used in each analysis. As this method determines the probability of ancestry from a diploid progenitor population relative to a non-admixed *C. bursa-pastoris* subgenome population, we defined regions of the subgenomes as putatively introgressed if the probability of ancestry from the progenitor diploid was greater than 50%. To check for reproductive barriers between *C. bursa-pastoris* and its diploid relatives, we performed artificial crosses. The crosses were made in both directions using *C. bursa-pastoris* as a mother plant and as a pollen donor. Each cross was replicated at least three times and each biological replicate consisted of 5 or more siliques. The details are provided in the Appendix S1.

### Selection tests

To search for selective sweeps, we used SweepFinder2 (74). SweepFinder2 was run on the data-set that besides polarized SNPs also included fixed derived alleles. This enables accounting for variation in mutation rate along the genome and increases power to detect sweeps (75). The critical composite likelihood ratio (CLR) values were determined using a 1% cut-off of the CLR values estimated in 100 simulations under a standard neutral model. The simulations were performed with *fastsimcoal2* (76). We assumed a mutation rate of 7 × 10^-9^ per site per generation, the population effective sizes for every population and subgenome were inferred from the *θ* values approximated by genetic diversity (*π*), and the average recombination rate was estimated using LDhelmet v1.7 (77). In addition, we estimated the ratio between nucleotide diversity at 0-fold (*π*_0_) and 4-fold degenerate sites (*π*_4_) in 5-6 samples with the lowest amount of missing data in each group. The details of the data used to estimate *π*_0_/*π*_4_ are provided in Fig. S5.

We also tested if the detected sweep regions were not the result of introgression or genome conversion. We compared the absolute genetic distance (*Dxy*) of each sweep region between all the groups and if the distance was the closest to one of the parental species or the opposite subgenome, such regions were classified as introgression or conversion, respectively. To reduce the number of potential false positives, we removed pericentromeric regions and all regions with repetitive sequences as annotated in Slotte et al. (59). Sweep regions with less than 10Kb apart were joined together and treated as one region.

### Genetic load estimation

To identify differences in genetic load between populations of *C. bursa-pastoris* (as well as to assess the effect of selfing on accumulation of deleterious mutations), we classified mutations into tolerated and deleterious ones using SIFT4G (38). We built the SIFT4G *Capsella rubella* reference partition database and used it to annotate our SNPs dataset. Then we analyzed the frequencies of tolerated and deleterious mutations. We also verified this analysis by using *A. thaliana* SIFT4G database and annotating *C. bursa-pastoris* according to the alignment between the two species. This verification was performed to make sure that the observed results were not due to a reference bias, because *C. rubella* is closer to *C. grandiflora* than to *C. orientalis*. To get only the annotation of the mutations that occurred after speciation of *C. bursa-pastoris*, we polarized the mutations with the reconstructed ancestral sequences (see above) and analyzed only derived mutations. We verified this polarization by analyzing only species(subgenome)-specific mutation (e.g. mutations unique to *C. bursa-pastoris* Co subgenome,*C. bursa-pastoris* Cg subgenome, *C. orientalis, C. grandiflora*, and *C. rubella*) (Fig. S14). All the counts were presented relative to the total number of annotated sites to avoid bias caused by variation in missing data between samples. The means of the genetic load were compared using the generalized linear model as we did for the topology weighting except that here we used a quasibinomial distribution due to overdispersion.

### Homeolog-specific expression analyses

Mapping of RNA-Seq reads to the *C. rubella* reference genome was conducted similarly to the mapping of DNA data using Stampy v1.0.22 (60) with the substitution rate set to 0.025. Although potential PCR duplicates are usually not removed from RNA-Seq data, for the allele-specific expression analysis removing duplicates is recommended (78). We marked duplicates with Picard Tools 1.115 and did not use them during the genotyping and homeolog-specific expression assessment. Variants were called using *HaplotypeCaller* (GATK) with heterozygosity set to 0.015, and minimum Phred-scaled call confidence of 20.0, and minimum Phred-scaled emit confidence of 20.0 as recommended for RNA-Seq data in GATK Best Practices (62). Among the obtained polymorphic sites those that had *MQ < 30.00, QD < 2.00, FS > 30.000* were filtered out. Calls with coverage of fewer than 10 reads were also excluded. Alleles counting was carried out using *ASEReadCounter* from GATK.

Homeolog-specific expression was assessed within the statistical framework developed by Skelly et al. (39). This framework uses a Markov chain Monte Carlo (MCMC) method for parameter estimation and incorporates information from both RNA and DNA data to exclude highly biased SNPs and calibrate for the noise in read counts due to statistical sampling and technical variability. First, we used DNA data to identify and remove SNPs that strongly deviated from the 0.5 mapping ratio. Second, we estimated the variation in allele counts using unbiased SNPs in the DNA data. Next, we fitted an RNA model using parameter estimated from DNA data in the previous step. Finally, we calculated a Bayesian analog of false discovery rate (FDR) with a posterior probability of homeologue specific expression (HSE) > 0.99 and defined genes with significant HSE given the estimated FDR. All inferences were performed using 200,000 MCMC iterations with bum-in of 20,000 and thin interval of 100. Each model was run three times with different starting parameters to verify convergence.

To test for differences between populations of *C. bursapastoris*, we analyzed phased expression data as was done with unphased data in Kryvokhyzha et al. (32). We tested differences between populations in two ways: each subgenome was processed individually in *edgeR*, and both subgenomes were analyzed together as proportional data by fitting a generalized linear model. In addition, we performed correction for genetic population structure by fitting generalized linear mixed models (see Appendix S2).

## DATA ACCESS

The sources of the data obtained from previous studies are provided in the Material and Methods. DNA sequences data generated for 21 accessions in this study is submitted to the NCBI database under the Sequence Read Archive number SRP126886. Both phased and unphased genotype data, phylogenetic trees, reconstructed ancestral sequences, estimates of *π* and *Dxy* with sliding window approach, results of PSMC and SMC++, SIFT annotations, CLR estimates of *sweepFinder2*, HAPMIX output, homeologue-specific gene expression values, and R scripts are deposited to the Dryad Digital Repository doi: XXXXXXX.

## AUTHOR CONTRIBUTIONS

DK and AC phased the data and performed selection tests. DK carried out phylogenetic and gene expression analyses. DK, MCE, TD, NT, TVK analyzed genetic load. JC performed demographic analyses and analyzed nucleotide diversity in the coding part of the genome. DK and MG did the crosses. DK, JK, and SG performed tests for introgression. DK and ML drafted the article with inputs from all other authors. JRS, UL, SG, SIW and ML supervised the project.

## ACKNOWLEDGEMENTS

We thank Clément Lafon-Placette and Mohammad Foteh Ali for help with crosses, and Pascal Milesi and Ludovic Dutoit for discussion of the results. We are especially grateful to Luis Leal for detailed feedback on the manuscript. Most of the analyses were carried out at Uppsala Multidisciplinary Center for Advanced Computational Science (UPPMAX) under the project b2013191. This study was supported by grants from the Swedish Research Council (VR) and the Erik Philip-Sörensens Stiftelse to ML.

## Supplementary Information

**Table S1. Sequencing and phasing information.**

**Table S2. Nucleotide diversity *π* and absolute divergence *D_xy_* between different subgenomes of *C. bursa-pastoris* populations and its parental species.**

**Table S3. Multiple comparisons for the generalized linear model of the topology weighting of the Co subgenome of *C. bursa-pastoris* and *C. orientalis.***

**Table S4. Multiple comparisons for the generalized linear model of the topology weighting of the Cg subgenome of *C. bursa-pastoris* and *C. rubella.***

**Table S5. Multiple comparisons for the generalized linear model of the topology weighting of the Cg subgenome of *C. bursa-pastoris* and *C. grandiflora.***

**Table S6. Summary of the probabilities and proportions of introgression from *HAPMIX* analyses of each of the *C. bursa-pastoris* subgenomes.**

**Table S7. Multiple comparisons for the generalized linear model on the genetic load estimated with *C. rubella SIFT4G* database.**

**Table S8. Multiple comparisons for the generalized linear model on the genetic load estimated with *A. thaliana* SIFT4G database.**

**Table S9. Multiple comparisons for the generalized linear model testing for the expression difference between populations.**

**Table S10. Alternative combinations used in the ABBA-BABA tests.**

**Fig. S1. Frequency of monophyly of different groups.**

**Fig. S2. Population size histories of *C.bursa-pastoris* and its parental species estimated with *PSMC* and *SMC++.***

**Fig. S3. Topology weighting of the three populations of**

***C. bursa-pastoris, C. orientalis*, and *C. grandiflora.***

**Fig. S4. Bar plots of *HAPMIX* introgression probabilities for major scaffolds (chromosomes) of *C. bursapastoris.***

**Fig. S5. Nucleotide diversity (*π*) in the coding part of the genome.**

**Fig. S6. Divergence between sequences in the regions of identified selective sweeps.**

**Fig. S7. Levels of heterozygosity in the phased and unphased data.**

**Fig. S8. Correlations between the levels of expression in the genes with significant homeologue-specific expression.**

**Fig. S9. Correlations between the levels of expression in all assessed genes.**

**Fig. S10. Distribution of *GATK* annotation scores.**

**Fig. S11. Example of the distribution of phasing states of haplotype blocks emitted by *HapCUT.***

**Fig. S12. Tree presumably reflecting the true history of the populations of *C. bursa-pastoris* and parental species.**

**Fig. S13. Maximum ikelihood phylogenetic tree of two subgenomes of *C. bursa-pastoris* and its parental species.**

**Fig. S14. Genetic load in the subgenomes of *C. bursapastoris* and its parental species for species-specific mutations.**

**Appendix S1. Artificial crosses**

**Appendix S2. Differential gene expression analysis between the populations of *C. bursa-pastoris* in the two subgenomes.**

